# Comprehensive mapping of local and diaspora scientists: a database and analysis of 63951 Greek scientists

**DOI:** 10.1101/2021.01.13.426588

**Authors:** John P.A. Ioannidis, Chara Koutsioumpa, Angeliki Vakka, Georgios Agoranos, Chrysanthi Mantsiou, Maria Kyriaki Drekolia, Nikos Avramidis, Despina G. Contopoulos-Ioannidis, Konstantinos Drosatos, Jeroen Baas

## Abstract

Research policy and planning for a given country may benefit from reliable data on both its scientific workforce as well as the diaspora of scientists for countries with substantial brain drain. Here we use a systematic approach using Scopus to generate a comprehensive country-level database of all scientists in Greece. Moreover, we expand that database to include also Greek diaspora scientists. The database that we have compiled includes 63951 scientists who have published at least 5 papers indexed in Scopus. Of those, 35116 have an affiliation in Greece. We validate the sensitivity and specificity of the database against different control sets of scientists. We also analyze the scientific disciplines of these scientists according to the Science Metrix classification (174 subfield disciplines) and provide detailed data on each of the 63951 scientists using multiple citation indicators and a composite thereof. These analyses demonstrate differential concentrations in specific subfields for the local versus the diaspora cohorts, as well as an advantage of the diaspora cohort in terms of citation indicators especially among top-impact researchers. The approach that we have taken can be applied to map also the scientific workforce of other countries and nations for evaluation, planning and policy purposes.

## INTRODUCTION

Construction of scientist databases can be useful tools for evaluation, planning and policy-making related to science. Effort to compile national databases of scientists with performance metrics, in particular citation indices, are sometimes undertaken by research assessment authorities.^1,2^ Often these efforts may not be sufficiently inclusive. For example, they may depend on non-systematic efforts where scientists voluntarily contribute information by themselves in order to be included. Moreover, citation metrics are difficult to standardize, especially when they are not calculated according to the same processes for all scientists, and when differences of scientific fields are insufficiently accounted for. Importantly, for many countries, brain drain is a major challenge for their scientific workforce.^3–5^ In these countries, planning and policy decisions would greatly benefit from mapping not only the disciplines and impact of scientists who still work in the country but also of those who have emigrated elsewhere. First generation emigrating scientists and often even second and higher generation emigrants may often still be interested in engaging with the scientific workforce of their country of origin, thus contributing valuable expertise. Countries with strong diaspora may benefit from the skills of diaspora scientists. <u>Scientific diaspora can be useful for both the mobile scientists^6–8^ and the countries involved at both ends as it can constitute a modern tool of scientific diplomacy and cooperation between the two countries.</u>^9,10^

Here we demonstrate how a large scale, standardized approach can be used to create an inclusive, comprehensive database of scientists in a specific nation. Moreover, we show how one can expand that database to include also scientists who have migrated to other countries. We focus our efforts on Greece and its national workforce and scientific diaspora. Greece is a country that has sustained over the years a very strong current of brain drain.^11–14^ Moreover, the country has been hit by a major economic crisis which has severely limited funding for research and development. Despite some improvements in recent years, funding remains highly suboptimal. Furthermore, scientists of Greek origin include many extremely influential scientists worldwide and past analyses suggest that there are many high-impact Greek scientists, both in Greece and abroad, who are leaders in their fields.^15^ Moreover, such previous work has suggested that the number of Greek scientists with substantial impact is much higher proportionally than the share of Greeks in the global population (10 million in Greece and perhaps another 3 million in the diaspora).^15^ Mapping Greek scientists in Greece and worldwide would be a valuable resource. The availability of comprehensive science publications databases such as Scopus and the fact that many first Greek names and the large majority of last Greek names tend to be highly specific for Greek descent allow creating a database of scientists of Greek origin. In this paper, we describe how we have constructed such a database and how we have examined its sensitivity and specificity in validation samples. We also present descriptive data for the entire database and for comparative evaluations of Greek origin scientists who have an affiliation in Greece and for those who have an affiliation in other countries. Our work may offer a template for similar scientist-mapping efforts on other countries.

## METHODS

### Eligibility criteria

We aimed to capture all scientists of Greek origin who have at least 5 published papers (articles, reviews, and conference proceedings). Eligible scientists were both those born in Greece and those born elsewhere (second or higher generation), but their family had a Greek origin. Scientists were eligible regardless of whether they had their current main affiliation in Greece or elsewhere. We excluded scientists who had fewer than 2 papers published after 1950.

In order to capture eligible scientists with an affiliation in Greece, we queried Scopus^16^ as of January 15, 2020 and identified all the last names that had at least one author ID (with any number of papers assigned) that included an affiliation in Greece. We found 70967 names where at least one author ID has an affiliation address in Greece. One researcher manually screened all of these names to identify those that seemed to be of Greek origin, allowing for inclusion of those who might be probable, to avoid losing potentially eligible names. A second researcher then examined the manual extraction and made amendments. Eventually, 57732 last names were retained.

We also screened manually the files of the top-100,000 most cited scientists based on a composite indicator that had been published previously.^17^ We used three different files of the top-cited scientists, each of which captured the top-100,000 including self-citations as well as the top-100,000 excluding self-citations based on career-long data in Scopus until the end of 2017 (http://dx.doi.org/10.17632/btchxktzyw.1#file-ad4249ac-f76f-4653-9e42-2dfebe5d9b01); based on citations received during a single calendar year, 2017 (http://dx.doi.org/10.17632/btchxktzyw.1#file-b9b8c85e-6914-4b1d-815e-55daefb64f5e); and based on career-long data until the end of 2018 (http://dx.doi.org/10.17632/btchxktzyw.1#file-bade950e-3343-43e7-896b-fb2069ba3481). These three files manually yielded 1044, 990, and 1013 eligible authors of Greek origin, with large overlap between the three lists.

In addition, two online sources of common Greek first names were screened manually starting from the most common ones until 86 first names were selected that were thought to be relatively specific for Greek origin people. For example, George is a common name in Greece, but it is not Greek-specific, i.e. the vast majority of people with first name George are not of Greek origin. Conversely, Georgios is highly Greek-specific.

At a next step, we retrieved from Scopus all author ID files with at least 5 papers (articles, reviews, or conference papers) where scientists had either a seemingly Greek-specific last name (any of the 57332 last names mentioned above, or any of the last names of highly-cited Greek scientists according to any of the three previously published lists) or a seemingly Greek-specific first name (any of the 86 mentioned above). Eventually, a total of 124656 author ID files were retrieved.

These 124656 files were manually screened, perusing information for each scientist including the first name, last name, country of listed affiliation, and institution of listed affiliation that could help identify if the scientist was of Greek origin or not. The availability of all scientists who shared one of the seemingly Greek-specific names along with country information allowed to identify whether any of these names were in fact not Greek-specific. Some last names occur identically both in Greeks and in some other nationality (e.g. Adam or Spinelli). In these cases, information on first name could help classify that individual if the first name was characteristically Greek. If the first name did not help to differentiate in this regard, the country information was used to arbitrate. The site https://forebears.io/ was consulted also in ambiguous cases, since it shows the relative frequency of surnames and names across different countries.

Of 124656 author files, it was concluded that 62837 were very likely Greek. We listed alphabetically by last names the 62837 authors and recorded additional first names that seemed to be Greek-specific. By screening 2,000 names at a time, it was found that relatively few new Greek-specific names were added after screening 8,000 authors and the incremental addition of eligible Greek origin authors would be limited by adding more first names. This process yielded a total of 370 seemingly Greek-specific first names and we then searched Scopus for all additional author IDs with these first names that had not been already captured in the 62837. These additional authors were then manually screened, and 1012 were deemed (based on their name and country information) to be eligible. The resulting database which comprised of 63849 author IDs was subjected to validation checks, as described below. Additions and deletions emerging during these validation checks and a final contribution by the authors of the present study of Greek scientists they knew of, who had not been captured, increased the final count by 102 to a final count of 63951 author IDs.

### Validation – sensitivity for capturing scientists of Greek origin who are in Greece

In order to evaluate the sensitivity of the compiled database in capturing scientists who work in Greece, we searched whether it had included scientists working at a university in Greece, the University of Thessaly. Scientists working in different universities and research institutions in Greece are not likely to have systematically different names, so one university is likely to provide a reasonably representative sample. We searched Google Scholar as the reference database since scientists need to enter their names and affiliations by themselves in creating a profile in Google Scholar. The 130 most-cited scientists with profiles and University of Thessaly affiliation in Google Scholar were screened and it was found that all of them (130/130) had been included in our compiled database. Therefore, the sensitivity was 100%, with binomial 95% confidence interval of 97.2%-100%.

### Validation – sensitivity for capturing scientists of Greek origin who are not in Greece

In order to evaluate the sensitivity of the compiled database in capturing scientists of Greek origin who do not work in Greece, we used two approaches.

First, we used a sample of scientists who had entered their names in a Linked In database of Greek biomedical scientists created by one of us (K.D.) for the World Hellenic Biomedical Association. We only considered names that had been entered by the scientists themselves, proving that they identified themselves as Greek; and we further limited the search to scientists who gave an address outside of Greece and who had a work title suggesting that they are faculty or other people in senior positions, as opposed to students. Of 42 such individuals, 34 were found to have at least 5 papers in Scopus. Of those 34, 26 were captured in the compiled database, for a sensitivity of 76.5% (95% confidence interval, 58.8% to 89.3%).

Second, we used the names of people listed in the Wikipedia entry on Greek Diaspora. These names are not necessarily of scientists, therefore we examined if each of the names would have been captured through either one of the Greek-specific last names or one of the first Greek-specific names that we had put together in order to compile our database of Greek scientists. For artists and other people who had acquired an artistic/stage name, we used their original name, since change to artistic/stage names would not apply for scientists. We excluded from the screening people born before 1900, as Greek names in the remote past may have been different. Eventually, 28 first-generation and 88 second- or later-generation Greeks were eligible for screening. 14/28 and 35/88 would have been captured by our last or first name searches, corresponding to sensitivity of 50% (95% confidence interval, 30.6% to 69.4%) and 39.8% (95% confidence interval, 29.4% to 50.8%), respectively.

### Validation – specificity for capturing scientists of Greek origin

In order to evaluate whether the compiled database might have captured any scientists who were not actually Greek, we randomly selected 100 of the 63849 author IDs. For each of them, we tried to find whether we could find their name written in Greek in the web. Of the 100, their Scopus affiliation was in Greece for 62, in Cyprus for 4, in other countries for 32, and for 2 authors we had no listed affiliation in Scopus. We could find their name written in Greek for all 100 authors. Therefore the specificity was 100% (95% confidence interval 96.4% to 100%).

### Evaluation of split author files

Some scientists in Scopus may have their published work split in 2 or more author ID files, and Scopus encourages authors to communicate directly with them to merge such split files. In order to assess how common this pattern might be in the compiled database of Greek authors, after listing the names alphabetically, every 100^th^ name was selected and assessed whether more than one author ID files may exist for that person in the database. Of 106 screened names, 9 (8.5%, 95% CI, 4.0 to 15.5%) had their work split in 2 files (n=8) or 3 files (n=1).

### Data included for each scientist in the database

From each author ID file included in the database, the following information is included based on data directly imported from Scopus on October 1, 2020 (when 7,983,030 author ID files with at least 5 papers [articles, reviews, or conference papers] were available in Scopus) and calculations that are the same to those performed for a recently published list of top-cited scientists:^18^ affiliation and country; publication year of earlier and latest Scopus-indexed publication, number of publications, number of publications in 1960-2020, 6 citation indicators and their composite (all indicators being presented both with and without self-citations), proportion of self-citations, ratio of citations to citing papers, ranking according to the composite indicator among all scientists worldwide with at least 5 papers, most common field of publications according to the 22-field Science Metrix classification, two most common sub-fields of publications according to the 174-subfield Science Metrix classification, and ranking according to the composite indicator among all the scientists in the same main (most common) Science Metrix subfield. For details on the Science Metrix classification see refs. 19 and 20. For authors where Scopus listed an affiliation but not a country, we tried to identify the country whenever it would be unambiguous based on the provided affiliation.

## RESULTS

### Main descriptive characteristics

Of the 63951 author ID files included in the final database, country of affiliation was available for 63174, and 35116 (55.6%) of them had their affiliation in Greece. Large shares of this cohort of scientists were also located in USA (n=9339, 14.8%), UK (n=6165, 9.8%), Germany (n=2083, 3.3%), Cyprus (n=2.7%), Australia (n=1155, 1.8%), France (n=1141, 1.8%%), Canada (n=1110, 1.8%), and Switzerland (n=994 n=1.6%) bus diaspora was worldwide (Figure 1).

**Figure 1:**
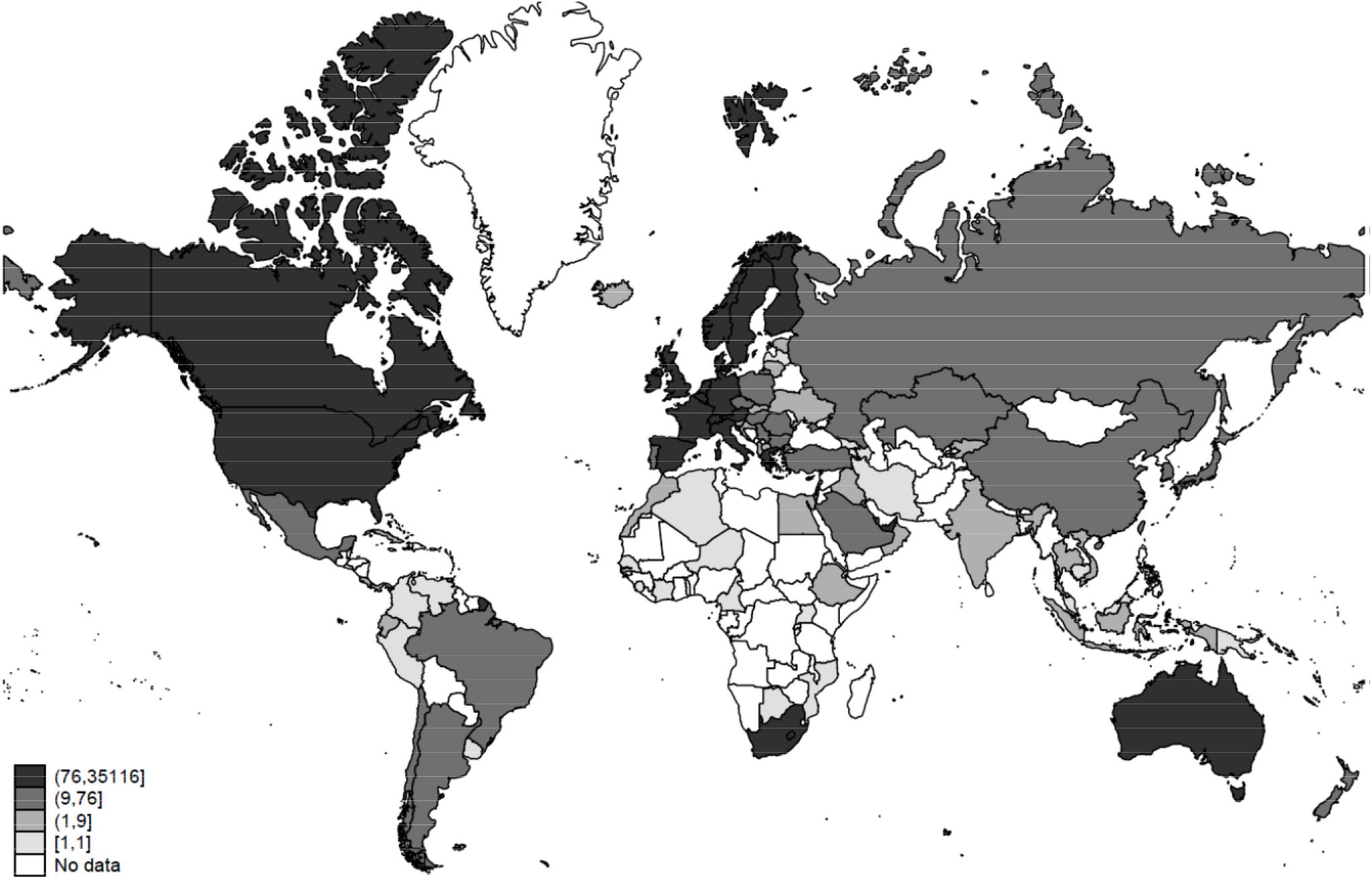
Worldwide distribution of scientists

12299 (19.2%) scientists had published their first Scopus-indexed paper after 2010 and 38248 (59.8%) had been recently active, publishing their last paper in 2018 or later. The median number of published papers was 13 (interquartile range, 8 to 31) and the median number of citations was 153 (interquartile range, 52 to 478).

As shown in Table 1, scientists with affiliation in Greece had an almost similar number of papers as scientists with affiliation outside of Greece, but they had substantially fewer citations, fewer papers that cited their work, and were placed on average in lower ranks compared with scientists with affiliation outside of Greece. Results were qualitatively similar regardless of whether self-citations were counted or excluded (Table 1). Scientists with affiliation outside of Greece tended to have younger publication ages (median for year of first publication 2004 versus 2002).

**Table 1.** Characteristics of scientists according to their country of current affiliation

| Characteristic, median (IQR) | Affiliation in Greece<br>N=35116 | Affiliation in other country<br>N=28058 |
| --- | --- | --- |
| Number of papers | 13 (7-30) | 14 (8-32) |
| Year of first paper | 2002 (1993-2008) | 2004 (1994-2010) |
| Year of most recent paper | 2019 (2013-2020) | 2019 (2014-2020) |
| Ranking across all science, thousands<br>(excluding self-citations) | 3505 (1691-5463) | 2812 (1265-4730) |
| Citations (excluding self-citations) | 139 (50-408) | 184 (59-609) |
| Citing papers (excluding self-citations) | 131 (48-375) | 169 (55-541) |
| Citations to citing papers ratio<br>(excluding self-citations) | 1.04 (1.00-1.11) | 1.07 (1.02-1.16) |
| Percentage of self-citations | 14 (7-25) | 15 (8-25) |
| Ranking across all science, thousands<br>(with self-citations) | 3508 (1681-5458) | 2801 (1259-4697) |
| Citations (with self-citations) | 169 (63-486) | 226 (76-730) |
| Ranking in main subfield (with self-citations) | 28711 (11338-61183) | 21378 (7500-48629) |
| Ranking in main subfield (without self-citations) | 28693 (11312-61180) | 21324 (7535-48751) |
| Percentile in main subfield (with self-citations) | 46 (22-71) | 37 (16-61) |
| Percentile in main subfield (without self-citations) | 46 (22-71) | 37 (16-61) |

A total of 33956 scientists with affiliation in Greece and 26150 scientists with affiliation in other countries could be assigned to a main scientific subfield. Among scientists who were in the top 0.1% of their subfield, the vast majority (86%) of them had an affiliation outside of Greece rather than in Greece (96 versus 15). For the top 0.5%, the respective numbers were 348 versus 89, for the top 1% the respective numbers were 648 versus 250, and for the top 5% the respective numbers were 2438 versus 1724, always with strong preponderance of scientists who were not in Greece. Below the top 5%, there was more equilibrium between scientists with affiliation outside of Greece versus in Greece, with the respective numbers being 7842 versus 7807 for the top 20%.

### Scientific fields

As shown in Table 2, Greek scientists had different representation across the 174 main scientific subfields of the Science Metrix classification. For 25 subfields, scientists in Greece exceeded by more than 2:1 the scientists with affiliation outside of Greece (Anatomy & Morphology, Environmental Engineering, Respiratory System, Obstetrics & Reproductive Medicine, Cardiovascular System & Hematology, Veterinary Sciences, Medical Informatics, Oceanography, Microbiology, Urology & Nephrology, Gastroenterology & Hepatology, General Clinical Medicine, Food Science, Marine Biology & Hydrobiology, Plant Biology & Botany, Fisheries, Sport Sciences, Environmental Sciences, Agronomy & Agriculture, Dairy & Animal Science, Entomology, Ornithology, History of Science, Technology & Medicine, Horticulture, Folklore). Conversely, in 32 subfields, scientists outside of Greece exceeded by more than 2:1 the scientists with affiliation in Greece (Cultural Studies, Law, Criminology, Communication & Media Studies, Art Practice, History & Theory, Economic Theory, Automobile Design & Engineering, Development Studies, General Chemistry, Social Work, Developmental Biology, Philosophy, Experimental Psychology, History of Social Sciences, Behavioral, Science & Comparative Psychology, International Relations, Psychoanalysis, Social Sciences Methods, Aerospace & Aeronautics, Human Factors, Developmental & Child Psychology, Applied Ethics, Anthropology, General Psychology & Cognitive Sciences, Social Psychology, Religions & Theology, Physiology, Literary Studies, Music, Clinical Psychology, Optics, History).

**Table 2.** Number of scientists in each scientific subfield

| Scientific subfield | Country |  |  |  |
| --- | --- | --- | --- | --- |
|  | Greece | Other | Unknown | Total |
| Accounting | 20 | 32 | 0 | 52 |
| Acoustics | 112 | 154 | 0 | 266 |
| Aerospace & Aeronautics | 57 | 159 | 3 | 219 |
| Agricultural Economics & Policy | 30 | 27 | 0 | 57 |
| Agronomy & Agriculture | 211 | 54 | 0 | 265 |
| Allergy | 80 | 62 | 0 | 142 |
| Analytical Chemistry | 368 | 194 | 2 | 564 |
| Anatomy & Morphology | 35 | 17 | 0 | 52 |
| Anesthesiology | 130 | 81 | 0 | 211 |
| Anthropology | 16 | 40 | 0 | 56 |
| Applied Ethics | 7 | 18 | 1 | 26 |
| Applied Mathematics | 160 | 100 | 1 | 261 |
| Applied Physics | 632 | 586 | 0 | 1218 |
| Archaeology | 107 | 61 | 0 | 168 |
| Architecture | 5 | 7 | 0 | 12 |
| Art Practice, History & Theory | 1 | 6 | 0 | 7 |
| Arthritis & Rheumatology | 194 | 126 | 0 | 320 |
| Artificial Intelligence & Image Processing | 1685 | 1293 | 7 | 2985 |
| Astronomy & Astrophysics | 166 | 246 | 1 | 413 |
| Automobile Design & Engineering | 1 | 5 | 1 | 7 |
| Behavioral Science & Comparative Psychology | 6 | 20 | 0 | 26 |
| Biochemistry & Molecular Biology | 286 | 474 | 0 | 760 |
| Bioinformatics | 72 | 105 | 0 | 177 |
| Biomedical Engineering | 262 | 248 | 1 | 511 |
| Biophysics | 64 | 71 | 0 | 135 |
| Biotechnology | 181 | 128 | 0 | 309 |
| Building & Construction | 168 | 148 | 0 | 316 |
| Business & Management | 200 | 203 | 0 | 403 |
| Cardiovascular System & Hematology | 1671 | 768 | 10 | 2449 |
| Chemical Engineering | 220 | 236 | 0 | 456 |
| Chemical Physics | 200 | 234 | 1 | 435 |
| Civil Engineering | 249 | 251 | 2 | 502 |
| Classics | 39 | 49 | 1 | 89 |
| Clinical Psychology | 24 | 51 | 0 | 75 |
| Communication & Media Studies | 9 | 56 | 0 | 65 |
| Complementary & Alternative Medicine | 6 | 8 | 0 | 14 |
| Computation Theory & Mathematics | 123 | 119 | 1 | 243 |
| Computer Hardware & Architecture | 173 | 164 | 0 | 337 |
| Criminology | 7 | 45 | 0 | 52 |
| Cultural Studies | 0 | 11 | 0 | 11 |
| Dairy & Animal Science | 214 | 54 | 0 | 268 |
| Demography | 13 | 10 | 0 | 23 |
| Dentistry | 421 | 287 | 5 | 713 |
| Dermatology & Venereal Diseases | 209 | 150 | 1 | 360 |
| Design Practice & Management | 24 | 35 | 0 | 59 |
| Development Studies | 3 | 13 | 0 | 16 |
| Developmental & Child Psychology | 35 | 94 | 0 | 129 |
| Developmental Biology | 172 | 712 | 0 | 884 |
| Distributed Computing | 64 | 94 | 0 | 158 |
| Drama & Theater | 3 | 2 | 0 | 5 |
| Ecology | 119 | 72 | 1 | 192 |
| Econometrics | 8 | 13 | 0 | 21 |
| Economic Theory | 4 | 22 | 0 | 26 |
| Economics | 310 | 286 | 0 | 596 |
| Education | 456 | 329 | 0 | 785 |
| Electrical & Electronic Engineering | 293 | 267 | 0 | 560 |
| Emergency & Critical Care Medicine | 183 | 93 | 0 | 276 |
| Endocrinology & Metabolism | 577 | 342 | 9 | 928 |
| Energy | 869 | 591 | 7 | 1467 |
| Entomology | 195 | 48 | 0 | 243 |
| Environmental & Occupational Health | 14 | 25 | 0 | 39 |
| Environmental Engineering | 246 | 119 | 1 | 366 |
| Environmental Sciences | 534 | 163 | 0 | 697 |
| Epidemiology | 43 | 30 | 0 | 73 |
| Evolutionary Biology | 64 | 57 | 0 | 121 |
| Experimental Psychology | 39 | 138 | 1 | 178 |
| Family Studies | 4 | 3 | 0 | 7 |
| Finance | 95 | 159 | 0 | 254 |
| Fisheries | 146 | 47 | 0 | 193 |
| Fluids & Plasmas | 159 | 195 | 0 | 354 |
| Folklore | 2 | 0 | 0 | 2 |
| Food Science | 377 | 132 | 4 | 513 |
| Forestry | 80 | 44 | 0 | 124 |
| Gastroenterology & Hepatology | 661 | 252 | 2 | 915 |
| Gender Studies | 5 | 7 | 0 | 12 |
| General & Internal Medicine | 375 | 231 | 5 | 611 |
| General Chemistry | 12 | 51 | 0 | 63 |
| General Clinical Medicine | 102 | 38 | 0 | 140 |
| General Mathematics | 237 | 162 | 0 | 399 |
| General Physics | 131 | 151 | 0 | 282 |
| General Psychology & Cognitive Sciences | 2 | 5 | 0 | 7 |
| Genetics & Heredity | 96 | 174 | 0 | 270 |
| Geochemistry & Geophysics | 270 | 167 | 8 | 445 |
| Geography | 37 | 26 | 0 | 63 |
| Geological & Geomatics Engineering | 272 | 187 | 1 | 460 |
| Geology | 27 | 20 | 0 | 47 |
| Geriatrics | 26 | 32 | 0 | 58 |
| Gerontology | 17 | 17 | 0 | 34 |
| Health Policy & Services | 46 | 78 | 0 | 124 |
| History | 19 | 39 | 1 | 59 |
| History of Science, Technology & Medicine | 18 | 4 | 0 | 22 |
| History of Social Sciences | 2 | 7 | 0 | 9 |
| Horticulture | 35 | 6 | 0 | 41 |
| Human Factors | 28 | 77 | 0 | 105 |
| Immunology | 706 | 487 | 1 | 1194 |
| Industrial Engineering & Automation | 332 | 390 | 0 | 722 |
| Industrial Relations | 10 | 14 | 0 | 24 |
| Information & Library Sciences | 38 | 34 | 1 | 73 |
| Information Systems | 143 | 204 | 4 | 351 |
| Inorganic & Nuclear Chemistry | 250 | 128 | 1 | 379 |
| International Relations | 16 | 49 | 0 | 65 |
| Languages & Linguistics | 57 | 57 | 0 | 114 |
| Law | 12 | 82 | 2 | 96 |
| Legal & Forensic Medicine | 30 | 23 | 0 | 53 |
| Literary Studies | 12 | 27 | 0 | 39 |
| Logistics & Transportation | 173 | 155 | 1 | 329 |
| Marine Biology & Hydrobiology | 247 | 83 | 1 | 331 |
| Marketing | 56 | 75 | 0 | 131 |
| Materials | 390 | 280 | 1 | 671 |
| Mathematical Physics | 38 | 26 | 0 | 64 |
| Mechanical Engineering & Transports | 253 | 218 | 1 | 472 |
| Medical Informatics | 184 | 78 | 0 | 262 |
| Medicinal & Biomolecular Chemistry | 236 | 149 | 0 | 385 |
| Meteorology & Atmospheric Sciences | 306 | 223 | 0 | 529 |
| Microbiology | 896 | 367 | 4 | 1267 |
| Microscopy | 6 | 4 | 0 | 10 |
| Mining & Metallurgy | 50 | 28 | 2 | 80 |
| Music | 6 | 13 | 0 | 19 |
| Mycology & Parasitology | 56 | 38 | 1 | 95 |
| Nanoscience & Nanotechnology | 75 | 134 | 0 | 209 |
| Networking & Telecommunications | 1100 | 953 | 10 | 2063 |
| Neurology & Neurosurgery | 742 | 962 | 3 | 1707 |
| Nuclear & Particle Physics | 509 | 623 | 0 | 1132 |
| Nuclear Medicine & Medical Imaging | 575 | 377 | 1 | 953 |
| Numerical & Computational Mathematics | 92 | 51 | 0 | 143 |
| Nursing | 147 | 89 | 1 | 237 |
| Nutrition & Dietetics | 230 | 126 | 0 | 356 |
| Obstetrics & Reproductive Medicine | 636 | 303 | 1 | 940 |
| Oceanography | 102 | 43 | 0 | 145 |
| Oncology & Carcinogenesis | 1718 | 868 | 3 | 2589 |
| Operations Research | 138 | 118 | 0 | 256 |
| Ophthalmology & Optometry | 301 | 308 | 0 | 609 |
| Optics | 74 | 155 | 0 | 229 |
| Optoelectronics & Photonics | 258 | 319 | 1 | 578 |
| Organic Chemistry | 232 | 212 | 0 | 444 |
| Ornithology | 17 | 4 | 0 | 21 |
| Orthopedics | 461 | 348 | 1 | 810 |
| Otorhinolaryngology | 206 | 128 | 0 | 334 |
| Paleontology | 54 | 34 | 0 | 88 |
| Pathology | 123 | 70 | 1 | 194 |
| Pediatrics | 226 | 113 | 2 | 341 |
| Pharmacology & Pharmacy | 396 | 208 | 0 | 604 |
| Philosophy | 12 | 44 | 2 | 58 |
| Physical Chemistry | 172 | 86 | 4 | 262 |
| Physiology | 26 | 59 | 0 | 85 |
| Plant Biology & Botany | 498 | 161 | 0 | 659 |
| Political Science & Public Administration | 55 | 107 | 0 | 162 |
| Polymers | 282 | 222 | 1 | 505 |
| Psychiatry | 277 | 265 | 2 | 544 |
| Psychoanalysis | 7 | 21 | 3 | 31 |
| Public Health | 87 | 139 | 0 | 226 |
| Rehabilitation | 36 | 68 | 0 | 104 |
| Religions & Theology | 7 | 17 | 0 | 24 |
| Respiratory System | 534 | 255 | 1 | 790 |
| Science Studies | 22 | 29 | 0 | 51 |
| Social Psychology | 30 | 74 | 1 | 105 |
| Social Sciences Methods | 4 | 12 | 0 | 16 |
| Social Work | 5 | 21 | 0 | 26 |
| Sociology | 12 | 18 | 0 | 30 |
| Software Engineering | 66 | 126 | 0 | 192 |
| Speech-Language Pathology & Audiology | 37 | 66 | 0 | 103 |
| Sport Sciences | 331 | 104 | 0 | 435 |
| Sport, Leisure & Tourism | 44 | 75 | 0 | 119 |
| Statistics & Probability | 144 | 132 | 0 | 276 |
| Strategic, Defence & Security Studies | 160 | 98 | 1 | 259 |
| Substance Abuse | 18 | 36 | 0 | 54 |
| Surgery | 760 | 493 | 16 | 1269 |
| Toxicology | 114 | 81 | 1 | 196 |
| Tropical Medicine | 34 | 32 | 0 | 66 |
| Urban & Regional Planning | 66 | 51 | 0 | 117 |
| Urology & Nephrology | 549 | 223 | 1 | 773 |
| Veterinary Sciences | 164 | 71 | 2 | 237 |
| Virology | 85 | 154 | 1 | 240 |
| Zoology | 33 | 18 | 0 | 51 |

### Top-cited Greek scientists across different fields

32 Greek scientists were among the top-15 of their scientific subfield based on a citation indicator excluding self-citations (Table 3). Almost all of them (30/32, 94%) were listed by Scopus with an affiliation outside of Greece. Of the 32 scientists, information on place of birth could be found on 28 (except for Terzopoulos, Stamatakis, Pavlou, Argyropoulos); three were born in Cyprus (Nicolaides, Nicolaou, Kalogirou), three were born in the USA (Ioannidis, Joannopoulos, Alivisatos), one was born in the UK (Lyketsos), and the remaining 21 had been born in Greece. Of the 32, 18 had received their first degree from an institution in Greece.

**Table 3.**
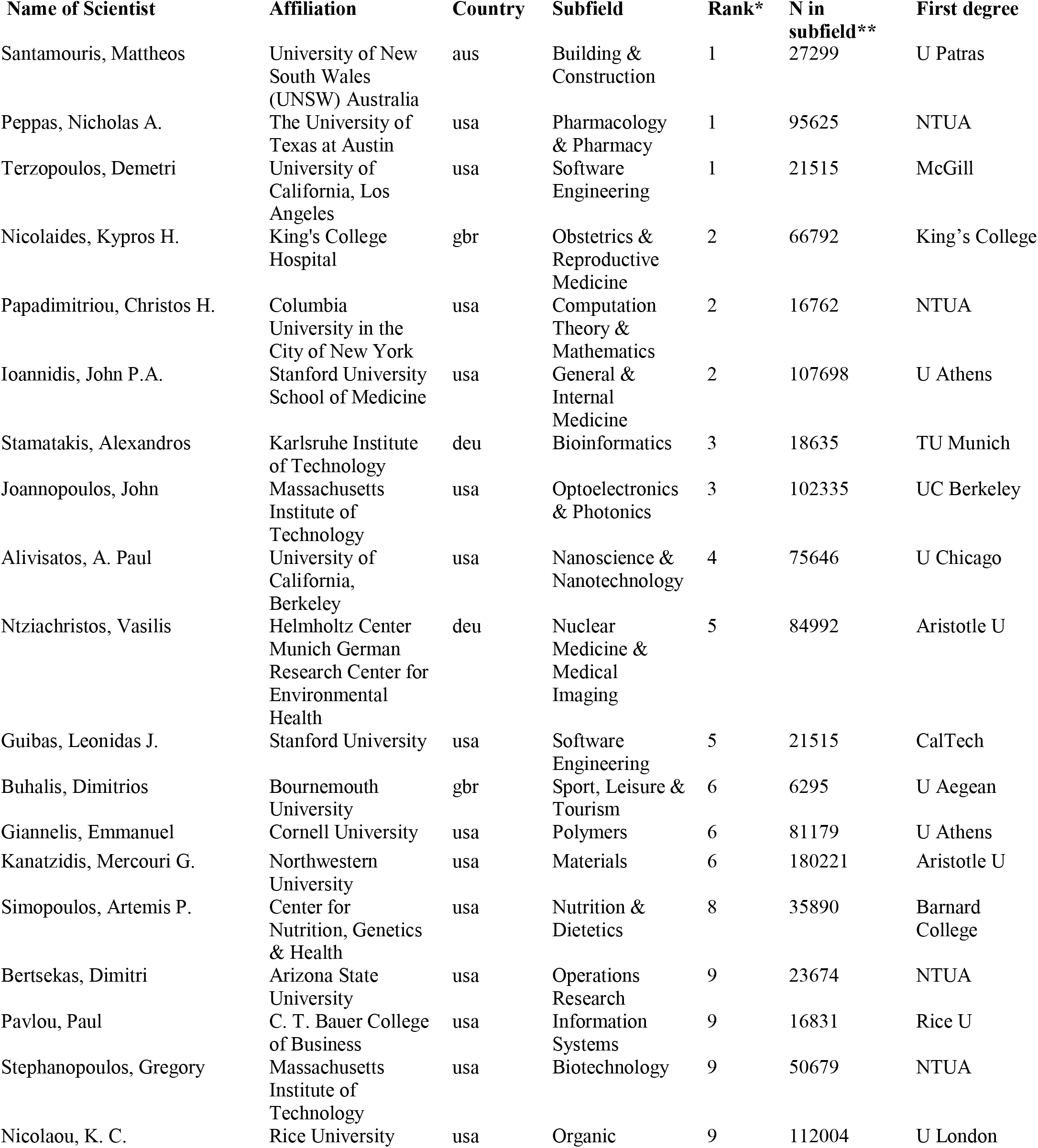
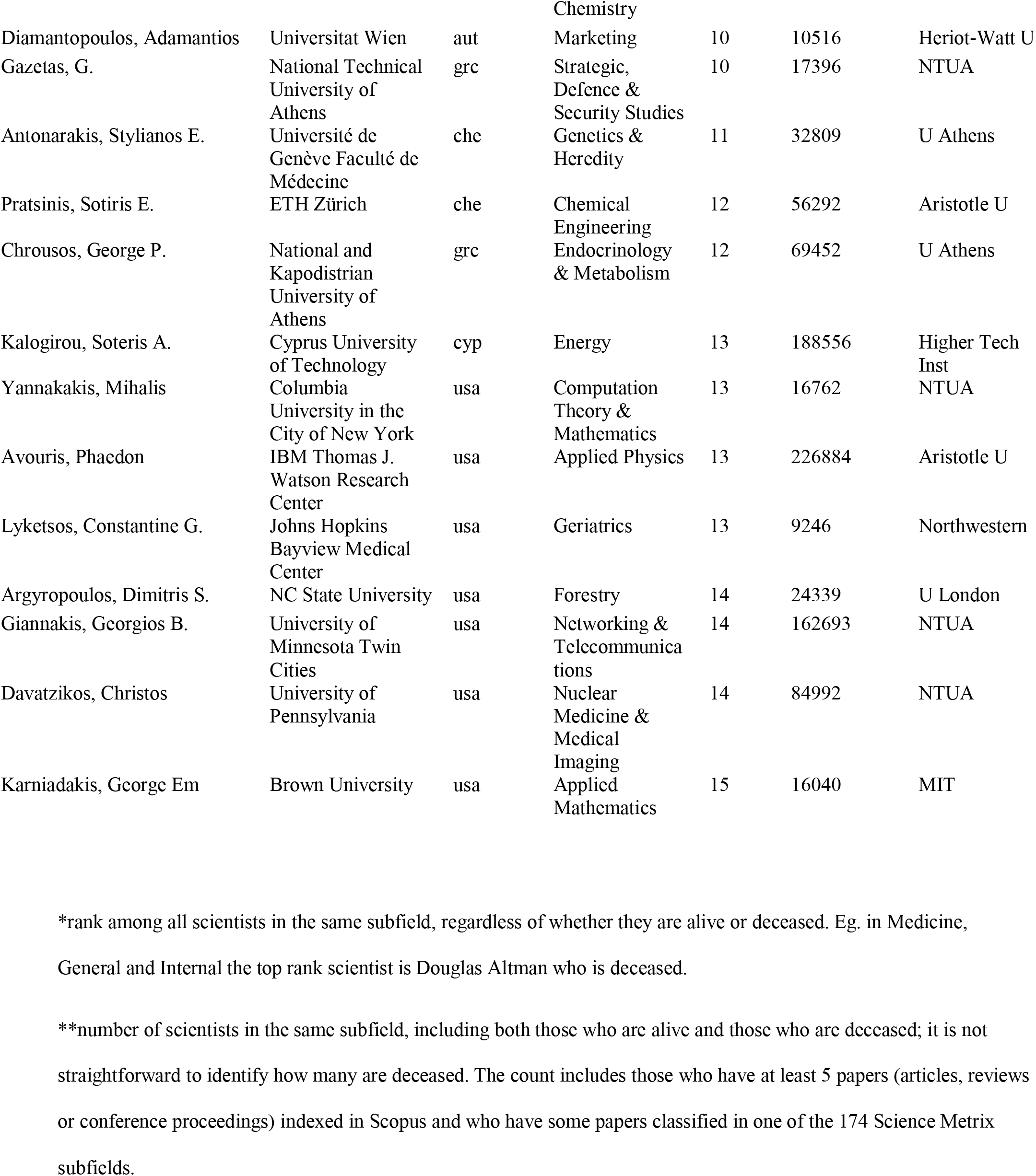
Scientists who are among the top-15 of their scientific subfield according to a composite citation indicator, excluding self-citations.

## DISCUSSION

We have created and validated a database of scientists of Greek origin that may be helpful for evaluation, planning, and research policy purposes. It may also serve as a template for similar efforts to be undertaken for other countries to map their scientific workforce. The iterative approach that we followed may also have special added value for countries that have sustained heavy brain drain and/or who have a substantial scientific diaspora.

The database includes close to 64000 author ID files representing scientists who have published at least 5 papers. Given that some scientists have their publications split in more than one file, the database probably includes close to 60000 unique scientists. Validation exercises suggest that it probably misses very few scientists who meet the productivity eligibility criteria and who have an affiliation in a Greek institution. Conversely, a more substantial proportion has been missed among those who have an affiliation in an institution outside of Greece. The estimate of the missingness in this regard varies according to different validation sets that we used. Based on scientists working abroad who on their own initiative offered to be included in a database of Greek scientists, about one in 4 scientists were missed with our approach. The percentage of missingness was higher based on a Wikipedia list of diasporeans, even more when extending beyond first generation emigrants. It is unavoidable that our approach would miss Greeks that acquire non-Greek names (upon 2^nd^ and subsequent generations and for people who change their names (e.g. through marriage, by making the name less foreign-sounding in their new country, or other reasons) and those who have a Greek last name that was not among the ones we searched for. Some of these individuals may still be captured if they possess a highly Greek-specific first name among the list of first names that we screened for. Therefore, even though scientists with an affiliation in Greece were a slight majority in the compiled database, Greek origin scientists with an affiliation outside of Greece would probably be the majority if all Greek origin scientists could have been retrieved. The total of Greek origin scientists meeting the productivity eligibility criteria may be in the range of 80,000-100,000 (∼1.00-1.25% of global total). Conversely, a few scientists included in the database by failed disambiguation. The validation process suggests that this situation is probably very uncommon.

The database reflects the large extent of general emigration from Greece as well as the massive brain drain that the country has sustained over the years, with accelerated rates in the last decade in conjunction with the economic crisis that hit Greece worse than almost any other highly developed country. We noted that the cohort of scientists with affiliation outside of Greece had on average younger publication ages, as revealed by the year of their first paper; half of them published their first paper in 2004 or more recently.

While citation indicators are quite high for the entire database averages, scientists with an affiliation outside of Greece have substantially stronger citation indicators and higher rankings in their fields compared with scientists with affiliation in Greece. The difference is more prominent among top-cited scientists, where 86% of the Greek origin scientists who are in the top 0.1% of their subfield are not in Greece. Similarly, almost all (94%) of the Greek origin scientists who are among the top-15 of their subfield are not in Greece. Of interest, the large majority of these extremely highly-cited scientists were born in Greece, and the majority also received their first degree in Greece. This further demonstrates the power of the brain drain process. At the same time, scientists who have remained in Greece still include large numbers placed at the top-20% of their subfield. Thus, the local scientific workforce still has considerable capacity for excellence.

The database includes scientists scattered across almost every scientific subfield. Scientists with an affiliation in Greece have stronger concentrations than those with affiliations outside of Greece in many fields of clinical medicine, several fields of biology, and agriculture/fisheries/forestry. Greece has one of the highest rates of physicians per population in the world, if not the highest.^21^ Many of them are engaged in research, authoring or co-authoring papers, since scientific publications are requested and appraised not only for academic track positions, but even for regular clinical positions in the national health system. The advantage is that these incentives create a large pool of physicians with exposure to research. The disadvantage is that much of this research may not be of high quality and these authors have no lasting commitment to research. The concentration in subfields of agriculture, fisheries, and biology is explained probably by the nature of the economy, although agriculture and related fields have shrunk in latest years. Conversely, there are several other fields where most scientists of Greek origin do not work in Greece. This pattern is particularly strong in the social and economic sciences and some cutting-edge biomedical sciences, such as developmental biology.

Some limitations need to be discussed. First, as we have already acknowledged, the database is still missing several Greek origin scientists, in particular among those living and working abroad. We encourage providing relevant information at www.drosatos.com/greekscientists to bring such cases into our attention. While it is impossible to update the database adding one more scientist at a time, collecting information on missing individuals may allow us to consider further optimized automated processes in the future.

Second, the constructed database was restricted to scientists with at least 5 full papers. In the entire Scopus database, roughly four-fifths of author ID files have fewer than 5 papers. Some of the author ID files with sparse papers may be split-off fragments of the publication corpus of authors represented by some larger file. Nevertheless, by extrapolation, the total number of Greek authors who have published at least one paper may be in the range of 250,000-500,000. The overwhelming majority of authors of 1-4 papers are not major contributors or leaders in the scientific enterprise. However, many young scientists in this group may become major contributors or leaders in the future. Therefore, follow-up updates would be useful to perform.

Third, errors (either splitting the same author in two or more author ID files or including some papers by two or more authors in the same file) and inaccuracies in affiliations are possible. Authors who recognize errors should contact directly Scopus to make these corrections in Scopus per se, so that they may be carried over in our database, with any potential future updates. The entire Scopus database currently has overall 98.1% precision (proportion of papers in an author ID file that belong to the author) and 94.4% recall (proportion of papers of an author included in the largest profile). Precision and recall may be even better for Greek-name authors, because Greek names are more rare and thus more specific than those of most other origins (e.g. disambiguation challenges for “Liu Wang” are greater than for “Yiannis Triantafyllou”). We found 8.5% of the authors in the database to have a split profile and, given that even when one profile carries the large majority of the author’s papers, recall probably substantially exceed 94.4% for our database.

Fourth, allocation of fields and subfields follows a well-established classification, but some scientists may have an almost equal number of papers in two or more fields, and the most common one may not fully capture their expertise. Their ranking would have been different, had they been classified in a different subfield. Moreover, even within the same subfield, there are granular subsections with different citation densities.

Fifth, allocation of affiliation and country is performed automatically by Scopus picking just one affiliation from the most recent papers of each author. Some authors have multiple current affiliations, and some may have changed their affiliation recently. Again, we encourage authors who want to change their listed affiliation to communicate directly with Scopus. Misclassification may affect some authors in their classification as being in Greece versus outside of Greece. However, it would have been extremely difficult to curate affiliations manually and it is impossible for an outsider to know which of many affiliations an author may prefer.

Finally, all citation metrics have limitations and their use should be done with caution and not as absolute indicators.^22–24^ We made no effort to assess the quality of the published works. Some authors may rank high, but may have other reasons for concern, e.g. retracted papers, or implausibly high self-citation metrics or evidence for citation farms. These need to be carefully scrutinized on a case-by-case basis.

Acknowledging these caveats, the compiled database offers a tool that may be useful for both research and policy purposes. For a country that is trying to recover from a lengthy economic crisis and a superimposed crisis from the recent COVID-19 pandemic, realization of its scientific potential, deceleration and reversal of the brain drain and informed decision-making in the interface of science and society may offer substantial added value. Brain drain and diaspora does not need to have negative consequences for the home country; mapping of the scientific workforce and diaspora may help to maximize positive impact.^9,10,25^

We also hope that the iterative approach used here may be applied also to map the scientific workforce and scientific diaspora of other countries/nations as well. Scopus data can readily identify scientists with affiliation in a given country. In the case of Greece, where few scientists immigrate to from other countries, almost all scientists with affiliation in Greece have Greek names. This would not be true for countries that attract many scientists from other countries, but usually it is more important to map the entire scientific workforce rather than just native scientists. The ability to map the diaspora of different countries depends on whether there are many first and last names that are country-specific. Specificity may vary substantially across countries and careful validation and cross-checking procedures should be applied accordingly.

## Contributions

JPAI had the original idea which was further elaborated with KD and JB; all authors collected and curated data; JPAI, DGC-I and JB performed analyses; all authors interpreted the data; JPAI wrote the manuscript which was critically revised by all authors; all authors approved the final paper.

Data: All the data on the 63951 scientists are available in public view in Mendeley at https://doi.org/10.17632/zbyctscmbn.1

